# Mammalian-unique eIF4E2 maintains GSK3β proline kinase activity to resist senescence against hypoxia

**DOI:** 10.1101/2020.10.20.346569

**Authors:** Min zhang, Lei Sun, Dong He, Jian Chen, Zhiqiang Dong, Huiting Liang, Yu Cao, Bingcheng Cai, He Yang

## Abstract

Cellular senescence is a stable state of cell cycle arrest elicited by various stresses. Hypoxia modulates senescence, but its consequences and implications in living organisms remains unknown. Here we identified the eIF4E2-GSK3β pathway regulated by hypoxia to maintain p53 proline-directed phosphorylation (S/T-P) to prevent senescence. We previously knew that GSK3β activates p53 translation through phosphorylation of RBM38 Ser195 (-Pro196). Unexpectedly, eIF4E2 directly binds to GSK3β via a conserved motif, mediating Ser195 phosphorylation. Phosphoproteomics revealed that eIF4E2-GSK3β specifically regulates proline-directed phosphorylation. Peptide e2-I or G3-I that disrupts this pathway dephosphorylates p53 at multiple S/T-P, which accelerate senescence by transcriptional suppressing TOPBP1 and TRX1. Consistently, peptides induce liver senescence that is rescued by TOPBP1 expression, and mediate senescence-dependent tumor regression. Furthermore, hypoxia inhibits eIF4E2-GSK3β. Inspiringly, eIF4E2-GSK3β is unique to mammals, which maintains mice viability and prevents liver senescence against physiological hypoxia. Interestingly, this mammalian eIF4E2 protects heart of zebrafish against hypoxia. Together, we identified a mammalian -unique eIF4E2-GSK3β pathway preventing senescence and guarding against hypoxia in vivo.

## Introduction

Senescence is a stable cell-cycle arrest which limits tissue damage and tumorigenesis by excluding damaged cells (Chan & Narita, 2019; He & Sharpless, 2017). However, senescence paradoxically promotes cancer through senescence-associated secretory phenotype (SASP), which is a highly dynamic process that negatively affects the microenvironment (Lee & Schmitt, 2019). For similar reasons, senescence is also a potential contributor to aging or aging-related disease by profoundly affecting tissue homeostasis (Chan & Narita, 2019; He & Sharpless, 2017; Lee & Schmitt, 2019). Various cellular stresses promote senescence characterized by senescence-associated β-galactosidase activity (SA-β-gal), associated with similar phenotype (Chan & Narita, 2019; He & Sharpless, 2017). Hypoxia, one of shared features of cancer or aging, can avert stress-induced senescence in cultured cells (Leontieva et al, 2012; Welford & Giaccia, 2011). However, hypoxia can negatively or positively influence senescence in different physiological context, and the related mechanisms and their biological significance to living organisms are poorly understood (Childs et al, 2015; Mo et al, 2013; Xing et al, 2018).

The p53 transcription factor modulates senescence contributing to tumor suppression (Kastenhuber & Lowe, 2017; Vousden & Prives, 2009). p21, as a well-known transcriptional target of p53, is most critical to mediate senescence. Besides, p53 drive senescence via stimulating or inhibiting other targets (Xu et al, 2019a; Xu et al, 2019b). Phosphorylation is essential for p53 activation to trigger senescence (Jung et al, 2019; Kim et al, 2014). And p53 selectively activating or repressing target genes, which was extensively regulated by phosphorylation (Kastenhuber & Lowe, 2017). Differently, physiological p53 preserves tissues from senescence, but the underlying mechanism is unresolved (Kortlever et al, 2006; Rufini et al, 2013).

Glycogen synthase kinase-3β (GSK3β), as a serine/threonine kinase, is at the center of cellular signaling correlated with aging or related diseases (Beurel et al, 2015; Souder & Anderson, 2019). GSK3β deficiency contribute to senescent cell accumulation (Souder & Anderson, 2019). However, GSK3β appears to both promote and oppose senescence under different circumstances (Kim et al, 2013; Liu et al, 2008; Seo et al, 2008). GSK3β regulates p53 under stress conditions via phosphorylation or interaction (Qu et al, 2004; Turenne & Price, 2001; Watcharasit et al, 2002). GSK3β is constitutively active and is suppressed by phosphorylation at Ser9 (Beurel et al, 2015). Furtherly, protein complexes or subcellular distribution determines its activity (Beurel et al, 2015; He et al, 2019). GSK3β substrates require pre-phosphorylation (priming motif) or proline-directed, but how to distinguish them is poorly understood (Beurel et al, 2015; Wang et al, 2018).

The RNA-binding protein RBM38, a target of the p53, interacts with eukaryotic initiation factor 4E (eIF4E), thus selectively inhibiting p53 translation by preventing eIF4E from binding to mRNA (Zhang et al, 2011). GSK3β activates p53 translation by activating phosphorylation of RBM38 Ser195 (Ser195-Pro196), which impedes the RBM38-eIF4E interaction (Zhang et al, 2013). In this study, we found that RBM38 binds eIF4E2, a eIF4E-homologous protein (4E-HP). eIF4E2 is a translation initiation modulator and specifically regulate translation via interactions with RNA-binding proteins (Cho et al, 2005; Uniacke et al, 2012; Uniacke et al, 2014). Hypoxia stimulates a switch from eIF4E to eIF4E2 cap-dependent translation, which is essential for survive of cancer cells (Uniacke et al, 2014). Here, we unexpectedly found that eIF4E2 interacts with GSK3β through a conserved motif to regulate its proline-directed kinase activity to maintain multi-S/T-P phosphorylation of p53, which opposing senescence. Impressively, this eIF4E2-GSK3β pathway is unique to mammals, which resists senescence and protects mice or zebrafish against hypoxia.

## Results

### eIF4E2 regulates GSK3β proline kinase activity by directly interaction

We performed co-immunoprecipitation combined with mass spectrometry and found that eIF4E2 may be one of the partners of RBM38. GST pull-down assay was performed and showed that His-eIF4E2 physically interacted with GST-RBM38, and conversely, His-RBM38 binds to GST-eIF4E2 (Fig EV1A and B). In addition, we showed that endogenous eIF4E2 was detected in anti-RBM38 but not IgG immunoprecipitates (Fig EV1C). Conversely, RBM38 was detected in anti-eIF4E2 but not IgG immunoprecipitates (Fig EV1D).

To explore the function of RBM38 that depends on eIF4E2, we knock-down eIF4E2 by siRNA. Unexpectedly, knock-down of eIF4E2 significantly downregulated the phosphorylation of RBM38 S195 in HCT116 cells, but did not affect the protein expression level of RBM38 (Fig 1A). By using another siRNA targeting eIF4E2, the same results were observed in both A549 and MCF7 cells (Fig 1B). To ensure results, two antibodies were used to detect Rbm38 S195 phosphorylation. The antibody p-RBM38#1 was used before (Zhang et al, 2013; Zhang et al, 2018a), while p-RBM38#2 is more specific and used in most following studies without further indications (Fig EV1E).

**Figure 1.**
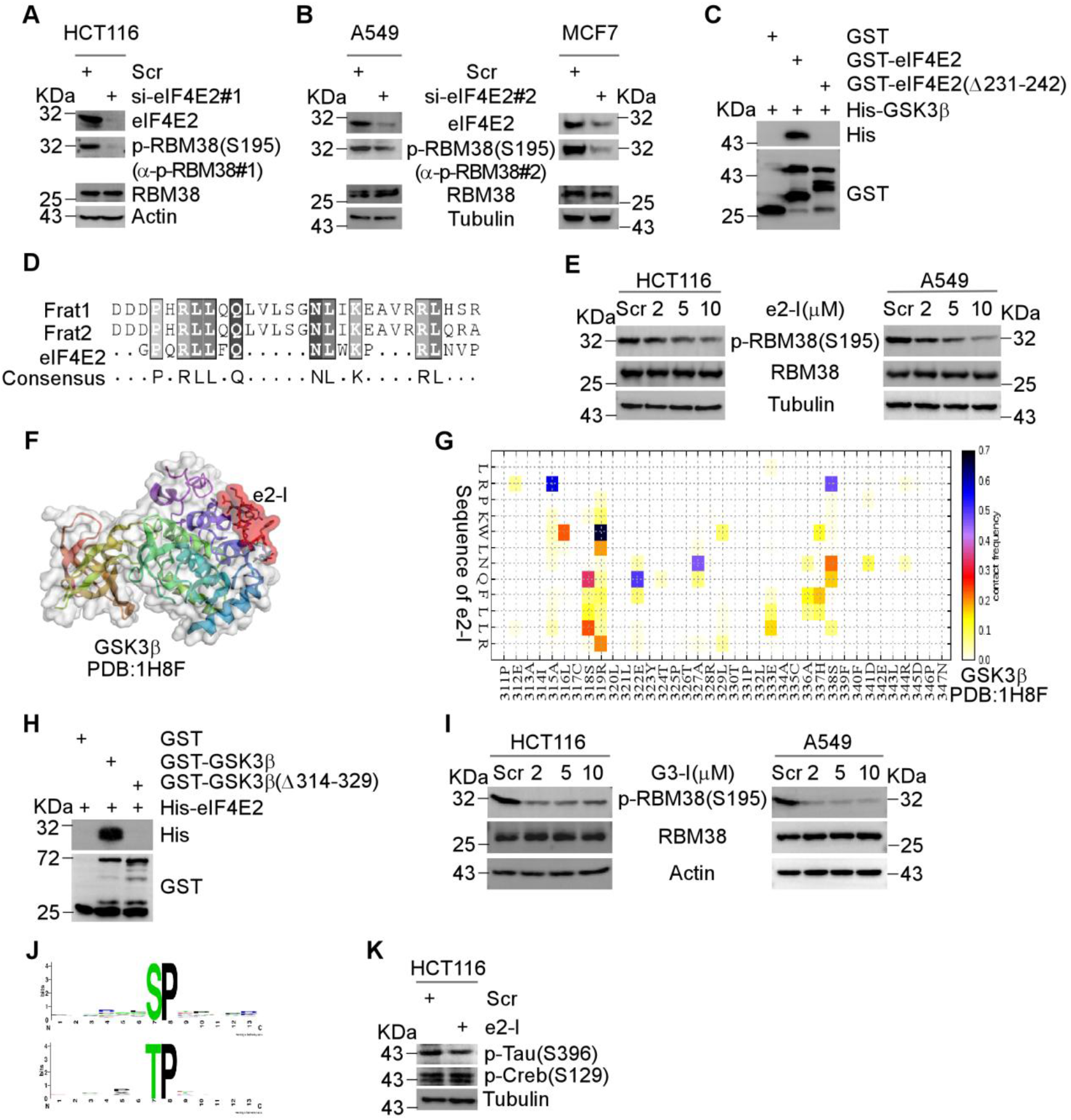
eIF4E2 regulates GSK3β proline kinase activity by directly interaction. **A** Depletion of eIF4E2 inhibits the phosphorylation of RBM38 S195 (Ser195-Pro). eIF4E2 siRNA#1 was transfected into HCT116 cells for 72 hours, followed by WB with indicated antibodies. Specifically, antibody p-RBM38#1 was used to detect p-RBM38(S195). **B** The experiment was performed as described in (A) except that eIF4E2 siRNA#2 was transfected into A549 or MCF7 cells. Specifically, antibody p-RBM38#2 was used to detect p-RBM38(S195). **C** Identifying the GSK3β binding motif of eIF4E2. GSK3β directly interacts with eIF4E2, but not with eIF4E2 mutant lacking amin acid from 231 to 242 (Δ231-242). GST-agarose beads bound GST-eIF4E2 and GST-eIF4E2(Δ231-242) or GST proteins were incubated with purified His-GSK3β. The elution was analyzed by WB using indicated antibodies. **D** The GSK3β binding motif of eIF4E2 or FRAT (Frat1 and Frat2) were aligned. **E** e2-I inhibits the phosphorylation of RBM38 S195. Cells were treated with different concentrations of e2-I (2, 5, 10μM) or scrambled e2-S (10μM) for 24 hours, followed by WB with indicated antibodies. **F** CABS-dock showed the binding of e2-I to GSK3β. The crystallographic structure of GSK3β (PDB ID: 1H8F) was used and the interaction interface was highlighted in red. **G** The contact diagram of GSK3β with e2-I, proposed by CABS-dock web server. **H** Identifying the eIF4E2 binding motif of GSK3β. GST-agarose beads bound GST-GSK3β and GST-GSK3β(Δ314-329) or GST proteins were incubated with purified His-eIF4E2, followed by WB with indicated antibodies. **I** G3-I inhibits the phosphorylation of RBM38 S195. **J** The most over-represented motif is proline-directed serine/threonine regulated by e2-I. Phosphorylation motifs were extracted by using Motif-X algorithm for phosphoproteomic data and threshold for significance was set to P< 0.000001. The logo was generated by using the Weblogo. **K** e2-I inhibits proline-directed phosphorylation. Tau S396 is followed by proline, while Creb S129 is within priming motif. Cells were treated with 5μM e2-I or e2-S for 24 hours and analyzed by WB.

Since eIF4E2 did not alter GSK3β expression or its Ser9 phosphorylation, eIF4E2 may regulate GSK3β by interaction. GST pull-down assays demonstrated the direct binding of GSK3β to eIF4E2 (Fig EV1F and G). And Immunoprecipitation (IP) assays showed the endogenous interaction between eIF4E2 and GSK3β (Fig EV1H and I). By mapping interactions, we found eIF4E2 mutant (Δ231-242), lacking amino acid from 231 to 242, does not bind to GSK3β (Fig 1C). Coincidentally, this sequence (R-L-L-F-Q-N-L-W-K-P-R-L) is highly conserved with a known GSK3β binding motif of FRAT (Fig 1D) (Freemantle et al, 2002). Corresponding to this motif, peptide e2-I fused with cell-penetrating peptide was synthesized, and scrambled peptide e2-S, designed online (www.mimotopes.com), was used as a control. As expected, e2-I competitively inhibited the binding of eIF4E2 to GSK3β in GST pull-down assay (Fig EV1J). Importantly, e2-I inhibited RBM38 S195 (Ser195-Pro196) phosphorylation in a dose-dependent manner (Fig 1E). As a control, e2-I did not affect the protein expression of RBM38 (Fig 1E), and e2-S had no any effect even at more extensive treatment (Fig EV1K).

CABS-dock, for protein–peptide molecular docking, showed e2-I binds to GSK3β (Blaszczyk et al, 2016) (Fig 1F). Consistent with CABS-dock contact map (Fig 1G), eIF4E2 interacts with full-length GSK3β, but not with its deletion mutant (Δ314-329) in GST-pull down assay (Fig 1H). Peptide G3-I, corresponding to this motif, inhibited eIF4E2-GSK3β binding (Fig EV1L) and consequently inhibited RBM38 S195 phosphorylation, compared to the scrambled peptide (Fig 1I).

Then, we performed iTRAQ-based quantitative phosphoproteomic analysis in cells treated with e2-I or e2-S. The inhibition of RBM38 phosphorylation was checked to ensure the effectiveness of peptide. Globally, we identified 1882 phosphosites belonging to 1059 phosphoproteins with alteration upon e2-I treatment. Interestingly, Motif-x analysis revealed the proline-directed serine/threonine (S/T-P) as the most representative motif (Fig 1J and Fig EV1M) (Schwartz & Gygi, 2005). In addition, proline located close to S/T (S/T-X-P, X is any amino acid) is also potential targeted sites (Fig EV1M). In contrast, no priming motif was found in these identified sites. For verification, e2-I inhibited the phosphorylation of Tau S396 (-Pro397), which has been identified in phosphoproteomics, but did not affect Creb S129 phosphorylation (with priming motif) (Fig 1K) (Leroy et al, 2010; Wang et al, 2010). Then, we performed sequence analysis on about 200 potential GSK3β binding proteins retrieved from BioGrid (Stark et al, 2006). Approximately 20 proteins contained a similar motif: R-L-L-*X*_(3-10)_-L-K, and peptides derived from FRAT, MACF1 and NINEIN can inhibit RBM38 S195 phosphorylation (Fig EV1N) (Freemantle et al, 2002; Hong et al, 2000; Ka et al, 2014).

### Inhibition of eIF4E2-GSK3β dephosphorylates p53 at multi-S/T-P

Inhibition of eIF4E2-GSK3β may suppress p53 translation mediated by RBM38 dephosphorylation (Zhang et al, 2013). By L-azidohomoalaine (AHA) labeling (Zhang et al, 2019b), we showed that the newly synthesized p53 was decreased upon e2-I treatment, compared to e2-S (Fig 2A, left panel). However, e2-I had little effect on the whole protein expression of p53 (Fig 2A, right panel). In fact, e2-I oppositely extends the half-life of p53 protein, from around 5-20 min to 60-90 min (Lane, 1997) (Fig 2B). Consistently, e2-I inhibited the expression of cytoplasmic p53, but activated nuclear p53 expression (Fig 2C).

**Figure 2.**
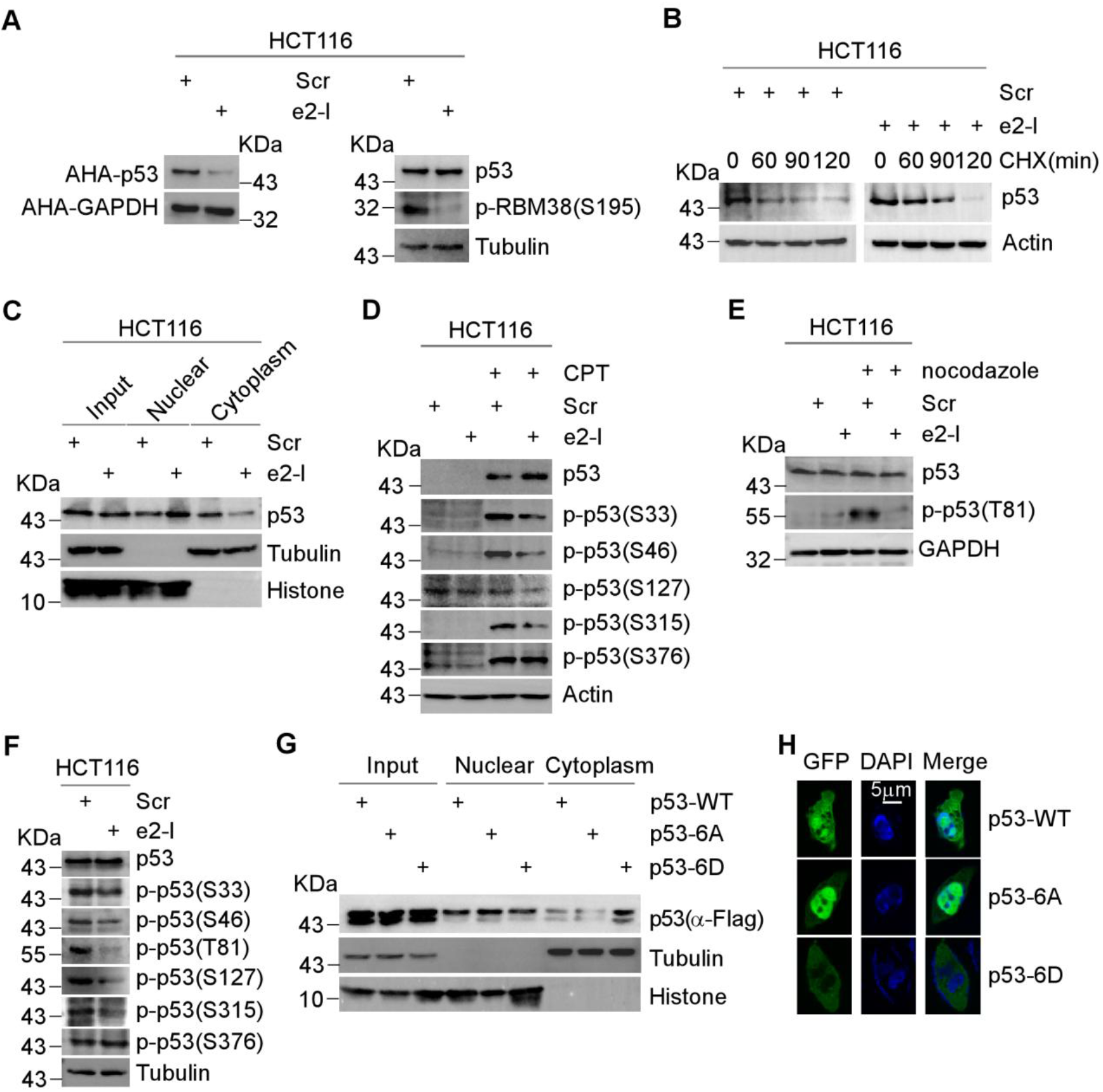
Inhibition of eIF4E2-GSK3β dephosphorylates p53 at multi-S/T-P. **A** e2-I inhibits p53 protein translation. Cells were treated with 5μM e2-I or scrambled e2-S for 24 hours. After cells were exposed to L-azidohomoalaine (AHA), cell lysates were incubated with the reaction buffer containing biotin/alkyne reagent. The biotin-alkyne-azide-modified protein complex was pulled down and analyzed by WB with indicated antibodies (left panel). The cell lysates before pull-down (input) were analyzed by WB with indicated antibodies (right panel). **B** e2-I extends the half-life of p53 protein. Cells were treated with peptide as in (A), then treated with or without cycloheximide (CHX, 0.1mg/mL) for up to 2 hours, followed by WB with indicated antibodies. **C** e2-I promotes nuclear localization of p53. Cells were treated with peptide as in (A), then the nuclear fraction was separated from the cytoplasmic fraction, followed by WB with antibodies against p53, Tubulin (markers for cytoplasmic) and Histone (markers for nuclear). **D** e2-I inhibits the CPT-induced phosphorylation of p53 at multi-S-P. Cells were treated with peptide as in (A), along with mock-treated or treated with 200 nM CPT for 24 hours, followed by WB with indicated antibodies. **E** e2-I inhibits the nocodazole-induced phosphorylation of p53 Thr81. The experiment was done as in (D), except that cells were mock-treated or treated with Nocodazole (50ng/ml). **F** e2-I inhibits multi-S/T-P phosphorylation of p53 at basal conditions. **G** Mutant p53-6A preferentially located in the nuclear. Vectors expressing FLAG tagged p53, p53-6A or p53-6D, were transfected into p53-null HCT116 cells for 48 hours. The nuclear and cytoplasmic fractions were separated, followed by WB with indicated antibodies. **H** Green fluorescent signals showed that mutant p53-6A is preferentially located in nuclear. Vectors expressing GFP-fused p53, p53-6A or p53-6D, were transfected into p53-null HCT116 cells for 24 hours. GFP fluorescence and DAPI staining were observed using confocal microscopy and the merged color coordinates of GFP and DAPI revealed the localization of p53.

We speculated that eIF4E2-GSK3β regulate S/T-P phosphorylation of p53 account for its increased stability (Fig EV2A). p53 phosphorylation at Ser33, Ser46, Ser315, were accumulated upon DNA damage elicited by camptothecin (Maclaine & Hupp, 2009) (Fig 2D). Activation of these S-P phosphorylation were blocked by e2-I (Fig 2D). Meantime, microtubule inhibitors nocodazole activated Thr81 phosphorylation (Buschmann et al, 2001), which was also inhibited by e2-I (Fig 2E). By using homemade antibody, we found Ser127 phosphorylation does not response to stress (Wei et al, 2003), and Thr150 phosphorylation remains undetectable. Notably, except Thr150, all basal S/T-P phosphorylation were suppressed by e2-I in resting cells (Fig 2F). For comparison, Ser376 phosphorylation, a known GSK3β substrate without proline-directed, was examined and showed no change (Fig 2D-F) (Qu et al, 2004). Similarly, G3-I inhibited p53 S/T-P phosphorylation in A549 cells under both stress (Fig EV2B and C) or basal conditions (Fig EV2D).

The nuclear p53 accumulation may be due to S/T-P dephosphorylation of p53, which subsequently preventing its cytoplasmic degradation. To test this, we generated vectors expressing p53 mutant 6A (with six S/T-P sites mutated to A-P, A is alanine) or 6D (mutated to D-P, D is aspartic acid), mimicking unphosphorylated or phosphorylated p53 at S/T-P, respectively. These vectors, as well as a control vector expressing wile-type p53, were introduced into p53-null HCT116 cells. We found that p53-6A is preferentially located in nucleus, whereas p53-6D is preferentially in cytoplasm (Fig 2G). GFP fluorescence fused with p53 also showed that 6A is majorly in nucleus, while 6D is in cytoplasm (Fig 2H). Together, we suggested that inhibition of eIF4E2-GSK3β leads to nuclear accumulation of p53 by dephosphorylating multi-S/T-P, which extends its half-life.

### Dephosphorylated p53 promotes senescence by repressing transcription

Next, we explored the function of eIF4E2-GSK3β pathway depending on p53. An increased SA-β-Gal (senescence-associated β-Galactosidase) staining was observed in e2-I treated HCT116 cells, but not in p53-null HCT116 cells (Fig 3A, upper panel). Similarly, e2-I promoted senescence in A549 cells (with wild-type p53), while had no significant effect in H1299 cells (p53-null) (Fig EV3A, upper panel). These two cell lines are both lung carcinoma cells with different endogenous p53 gene status. Consistently, e2-I activate the expression of senescence marker p21, depending on p53 (Fig 3A, lower panel and Fig EV3A, lower panel). Yet, e2-I did not induce apoptosis in above cells.

**Figure 3.**
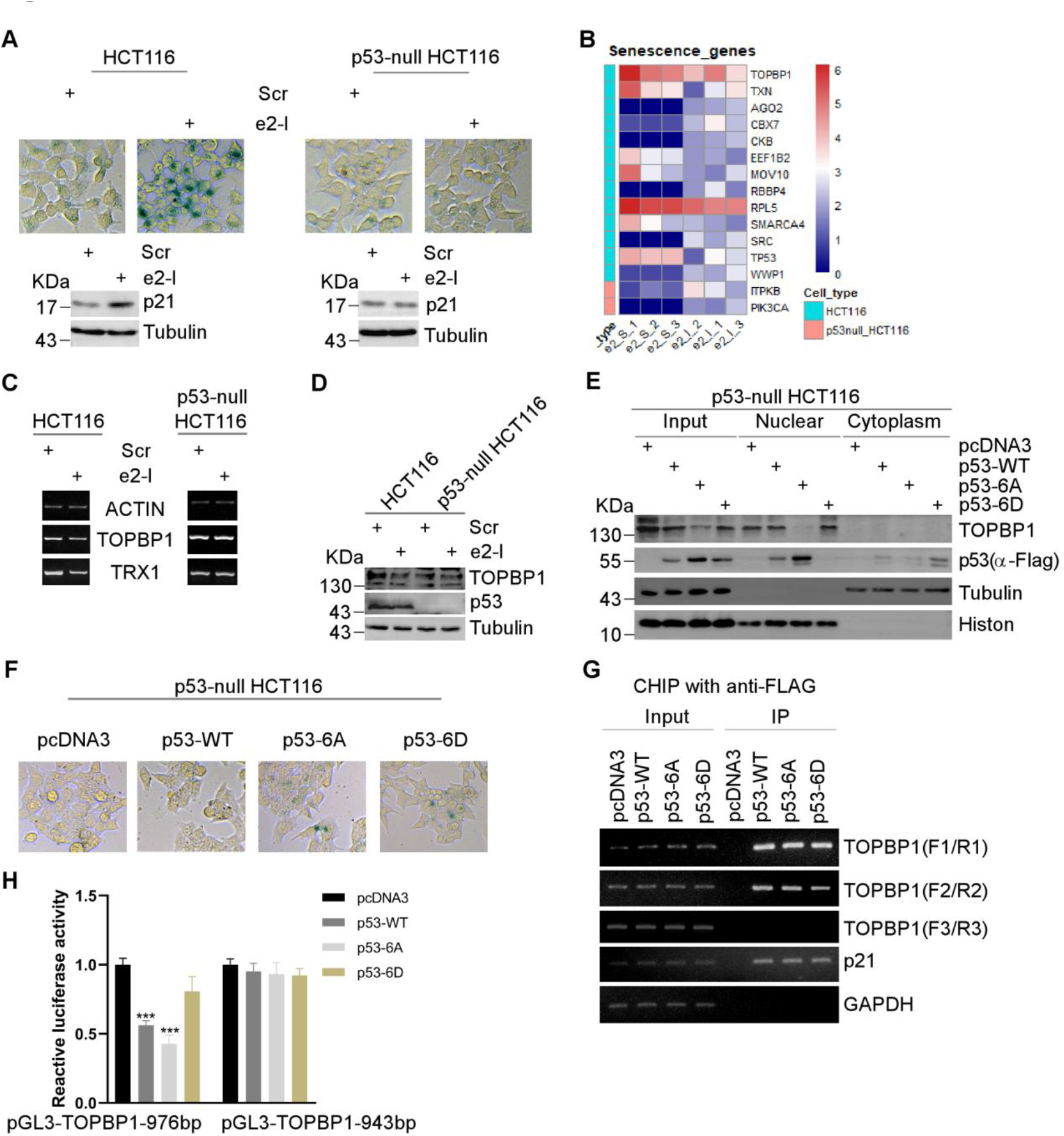
Dephosphorylated p53 promotes senescence by repressing transcription. **A** SA-β-Gal staining of HCT116 and p53-null HCT116 cells treated with 5μM e2-I or scrambled e2-S for 96 hours (upper panel). And cell extracts were subjected to WB with indicated antibodies (lower panel). **B** Heat map showed the differential expression genes associated with senescence in HCT116 and p53-null HCT116 cells, respectively. Differential expression genes (DEGs) were identified upon e2-I treatment by RNA-Seq. **C** e2-I inhibits mRNA expression of TOPBP1 or TRX1 depending on p53. HCT116 or HCT116-p53 null cells were treated with 5μM e2-I or scrambled e2-S for 24 hours. Total RNA was extracted and reverse transcriptase polymerase chain reaction (RT-PCR) was performed to check the mRNA expression. **D** e2-I downregulates the expression of TOPBP1 depending on p53. Cells were treated with peptide as in (c) and subjected to WB with indicated antibodies. **E** Mutant p53-6A suppresses the expression of TOPBP1. Vectors expressing FLAG tagged p53, p53-6A or p53-6D, were mock-transfected or transfected into p53-null HCT116 cells for 48 hours. Then, the nuclear fraction was separated from the cytoplasmic fraction, followed by WB with indicated antibodies. **F** Mutant p53-6A promotes senescence. SA-β-gal staining was performed in p53-null HCT116 cells, which were mock-transfected or transfected with vectors expressing p53, p53-6A or p53-6D, for 96 hours. **G** p53 associates with TOPBP1 promoter. FLAG-CHIP assay was performed in p53-null HCT116 cells, which were mock-transfected or transfected with vectors expressing FLAG-tagged p53, p53-6A or p53-6D, for 48 hours. The cell lysates were immunoprecipitated with anti-FLAG antibodies and analyzed by RT-PCR. **H** p53 directly inhibits TOPBP1 transcription activity. p53-null HCT116 cells were transfected with luciferase reporter constructs as indicated, along with mock-transfected or transfected with vectors expressing p53, p53-6A or p53-6D. Luciferase activity in cell lysates was measured and normalized by renilla activity using a dual-luciferase assay system. All data represent at least three independent experiments with similar results. p value <0.001 (***) vs control.

We performed transcriptome analysis to identify the underlying mechanism. GSEA assay showed WNT signaling pathway were significantly enriched in p53-null HCT116, but not in HCT116 upon e2-I treatment (Fig EV3B). And irrespective of p53 status, eIF4E2-GSK3β were significantly associated with neurodegenerative disease (Fig EV3C). Hence, eIF4E2-GSK3β pathway is highly related to WNT signaling, or neurodegenerative disease, which are known to be affected by GSK3β (Beurel et al, 2015).

Then, by referring to database of senescence-genes including CSGene (Zhao et al, 2016) or Cellage (genomics.senescence.info/cells/), 13 differentially expressed genes associated with senescence were found in HCT116 upon e2-I treatment, while only 2 in p53-null HCT116 (Fig 3B). Validation by western blotting and RT-PCR confirmed e2-I significantly inhibited TOPBP1 depending on p53 (Fig 3C and D). Consistently, p53-6A inhibited the expression of TOPBP1 in nuclear, whereas p53-wt or mutant 6D did not (Fig 3E). Moreover, e2-I or p53-6A repressed TRX1 (Fig 3C and Fig EV3D and EV3E). Coincidently, the inhibition of Topoisomerase II binding protein 1 (TOPBP1) and thioredoxin-1 (TRX1) promotes senescence by elevating p21 (Jeon et al, 2011; Young et al, 2010). Related, p53-6A expression promoted senescence (Fig 3F). Of note, p53-6D promoted senescence, after all it mimics phosphorylation activation and its targets requires further clarified (Fig 3F).

Using the Jaspar program, p53 responsive elements (REs) located approximately −5635 to −5620 and −5205 to −5187 bases upstream of the ATG transcription start site were found in promoter of TOPBP1 (Khan et al, 2018) (Fig EV3F). CHIP assays confirmed that p53 binds to TOPBP1 promoter region containing REs (Fig 3G, primer used for CHIP assay showed in Fig EV3F). Moreover, luciferase reporter assays showed p53-6A and p53-wt, but not 6D, inhibited the activity of the TOPBP1 pGL3-976bp construct that encompassing REs, while no effect on the pGL3-943bp construct lacking REs (Fig 3H and Fig EV3G). These data suggested p53 directly inhibits the expression of TOPBP1.

### Inhibition of eIF4E2-GSK3β promotes liver senescence rescued by expression

The physiological role of eIF4E2-GSK3β pathway was evaluated in both wild-type or *RBM38^S193A^* (corresponding to human S195) mutant mice (Fig EV4A). p53 were decreased in *RBM38^S193A^* MEF (mouse embryonic fibroblasts) compared to wild-type control (Fig 4A). e2-I had imperceptible effect on p53 expression in wild-type MEF, while it increased p53 expression in *RBM38^S193A^* MEF with more apparent senescence (Fig 4A and B).

**Figure 4.**
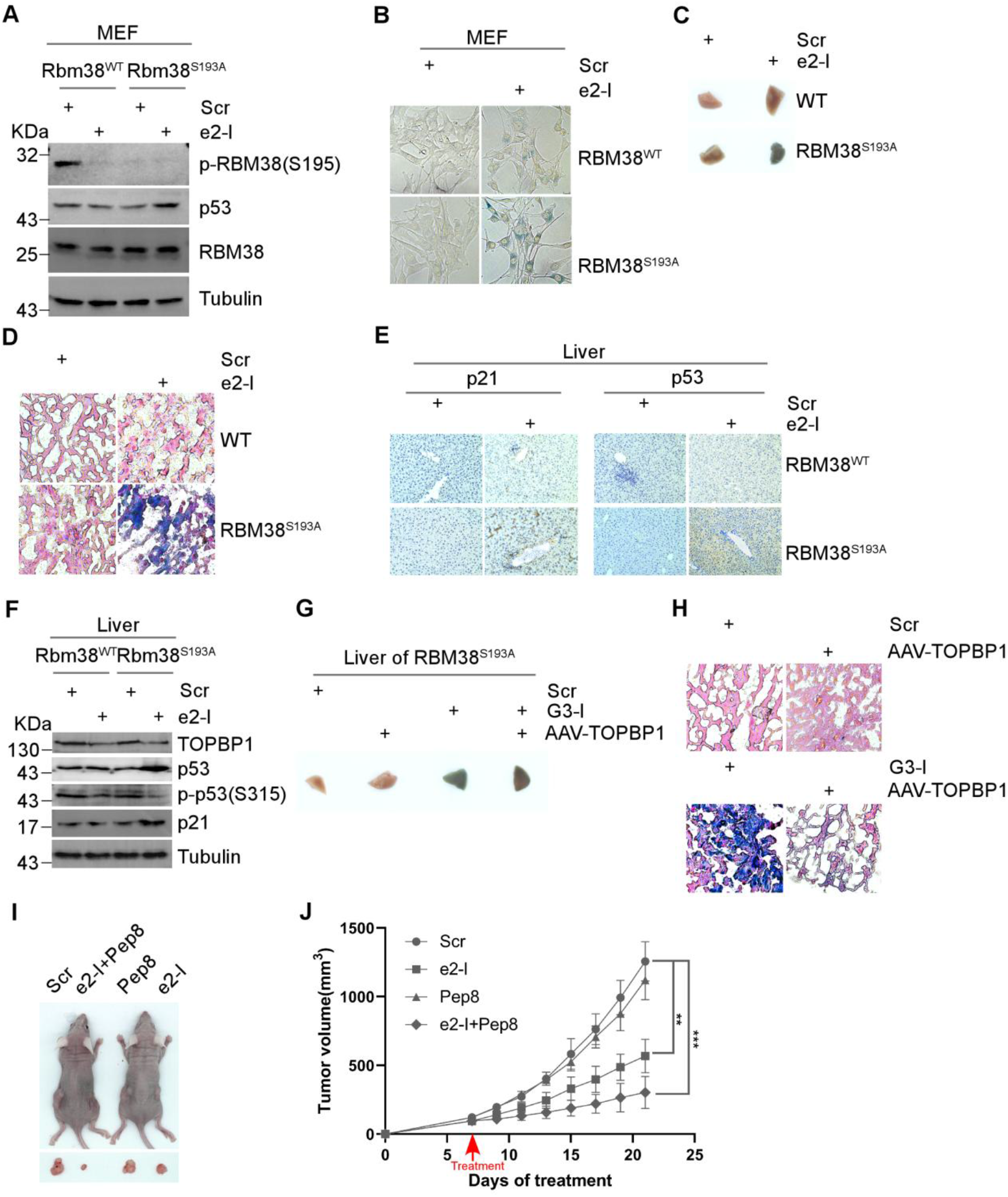
Inhibition of eIF4E2-GSK3β promotes liver senescence rescued by TOPBP1 expression. **A** e2-I promotes the expression of p53 in *RBM38^S193A^* MEF cells. MEF cells were treated with 5μM e2-I or the scrambled e2-S (5μM) for 24 hours and subjected to WB with indicated antibodies. **B** e2-I induced senescence in MEF cells. SA-β-Gal staining was performed in wild-type or *RBM38^S193A^* MEF cells treated with peptide for 96 hours. **C** e2-I promoted liver senescence. 35mg/kg e2-I or scrambled e2-S was intraperitoneal injected every two days, and on the 5^th^ day, SA-β-gal staining of liver tissue blocks were performed. **D** The SA-β-gal stained liver tissues shown in (C) were sectioned and then counterstained with nuclear fast red. **E** Immunohistochemical staining of liver tissue from experiment (C) for expression of p21 and p53. Scale bars, 50 μm. **F** Equal amounts of liver tissue extracts from experiment (C) were subjected to WB with indicated antibodies. **G** Expression of TOPBP1 rescued G3-I induced senescence. AAV-control or AAV-TOPBP1 (1×10^11^viral genomes in 100μl saline) were intravenous injected. 2 weeks later, mice were treated intraperitoneally with peptide G3-I as experiment (C), and on the 5^th^ day, SA-β-gal staining of liver tissue blocks were performed. **H** The SA-β-gal stained liver tissues shown in (G) were sectioned and then counterstained with nuclear fast red. **i**, Photographs of A549 xenografts after being excised from animal. A549 cells were injected into the 6-week-old male nude mice. After tumors reached a size > 100 mm^3^, intratumoral injections of 5μM e2-I were injected in combination with or without Pep8 (1μM) every other day for 14 days. **J** Xenograft tumor volume was measured every other day by using the following formula: ½ (Length x Width^2^), Statistical significance was determined by one-way analysis of variance, n=3 per group, and indicated by p value <0.001(***).

More important, intraperitoneal injection of e2-I strikingly promoted liver senescence in *RBM38^S193A^* mutant mice, compared to scrambled e2-S (Fig 4C and D). Meanwhile, e2-I hardly promoted liver senescence in wild-type mice (Fig 4C and D). Immunohistochemistry (IHC) showed that e2-I induced expression of p53 or p21 in liver of *RBM38^S193A^* mutant mice (Fig 4E). Western blotting further showed e2-I inhibited p53 S315 phosphorylation or TOPBP1 expression (Fig 4F). Furthermore, Interleukin-1-beta (IL-1β), one of SASP genes, were increased in *RBM38^S193A^* mutant mice upon e2-I treatment (Fig EV4B). ELISA measurements showed e2-I treatment increased secretion of IL-1β, IL-6, and IL-8, representing developed SASP (Fig EV4C). More important, G3-I or e2-I induced liver senescence in *RBM38^S193A^* mutant mice, which can be rescued by expression of TOPBP1 (Fig 4G and H and Fig EV4D and E). Consistent with above results in cultured cells, we suggested that loss of eIF4E2-GSK3β activity promotes senescence in vivo mediated by suppressing TOPBP1.

e2-I induced senescence may deter tumor progression. Xenograft tumor models showed that e2-I inhibited the growth of HCT116 xenografts, compared to scrambled e2-S. In contrast, e2-I has no effect on p53 null HCT116 xenografts (Fig EV4F and G). Then, the A549 xenograft model was used to further evaluate the antitumor activity of e2-I by intratumor injection after xenograft tumors reached 100 mm^3^ in size. e2-I reduces the growth of xenograft tumors compared to e2-S (Fig EV4H and I), and SA-β-Gal staining showed e2-I promotes senescence (Fig EV4J, left panel), accompanied by a decrease of TOPBP1 expression or an increase of p21 expression in xenograft tumors (Fig EV4J, right panel).

RBM38 Ser195 dephosphorylation may restrict the effect of e2-I by inhibiting p53, and a novel peptide Pep8 may relieve this restriction by disrupting RBM38-eIF4E binding (Lucchesi et al, 2019). Pep8, at a relative low concentration of 1μM, negligibly inhibited xenograft growth with no p53 induction. But impressively, combined use of Pep8 (1μM) and e2-I (5μM) substantially reduced the xenograft growth with activation of p53, compared to e2-I alone or Pep8 alone treatment (Fig 4I and J and Fig EV4K). These results suggested that reconciling the opposite effect of eIF4E2-GSK3β could be a promising cancer therapy strategy.

### Hypoxia inhibits eIF4E2-GSK3β activity by S-Nitrosylation

RBM38 S195 phosphorylation can be a representative of eIF4E2-GSK3β activity, and We found that hypoxia significantly inhibited the phosphorylation of RBM38 S195 (Fig 5A and Fig EV5A), suggesting that hypoxia restrains eIF4E2-GSK3β activity. As Fig 5B showed, eIF4E2 was detected in the anti-GSK3β immune complex under normal conditions, whereas hypoxia exposure (1% O_2_, 24 hours) significantly interfered with their interaction. To further test this, we performed a phase separation-based protein interaction reporter (SSPIER) analysis to visualize the dynamics of protein interaction (Chung et al, 2018; Zhang et al, 2018b). eIF4E2-GSK3β interaction leaded to the phase separation, prompting the formation of EGFP droplets, and hypoxia (1% O_2_) demolished EGFP droplets within 2 hours (Fig 5C).

**Figure 5.**
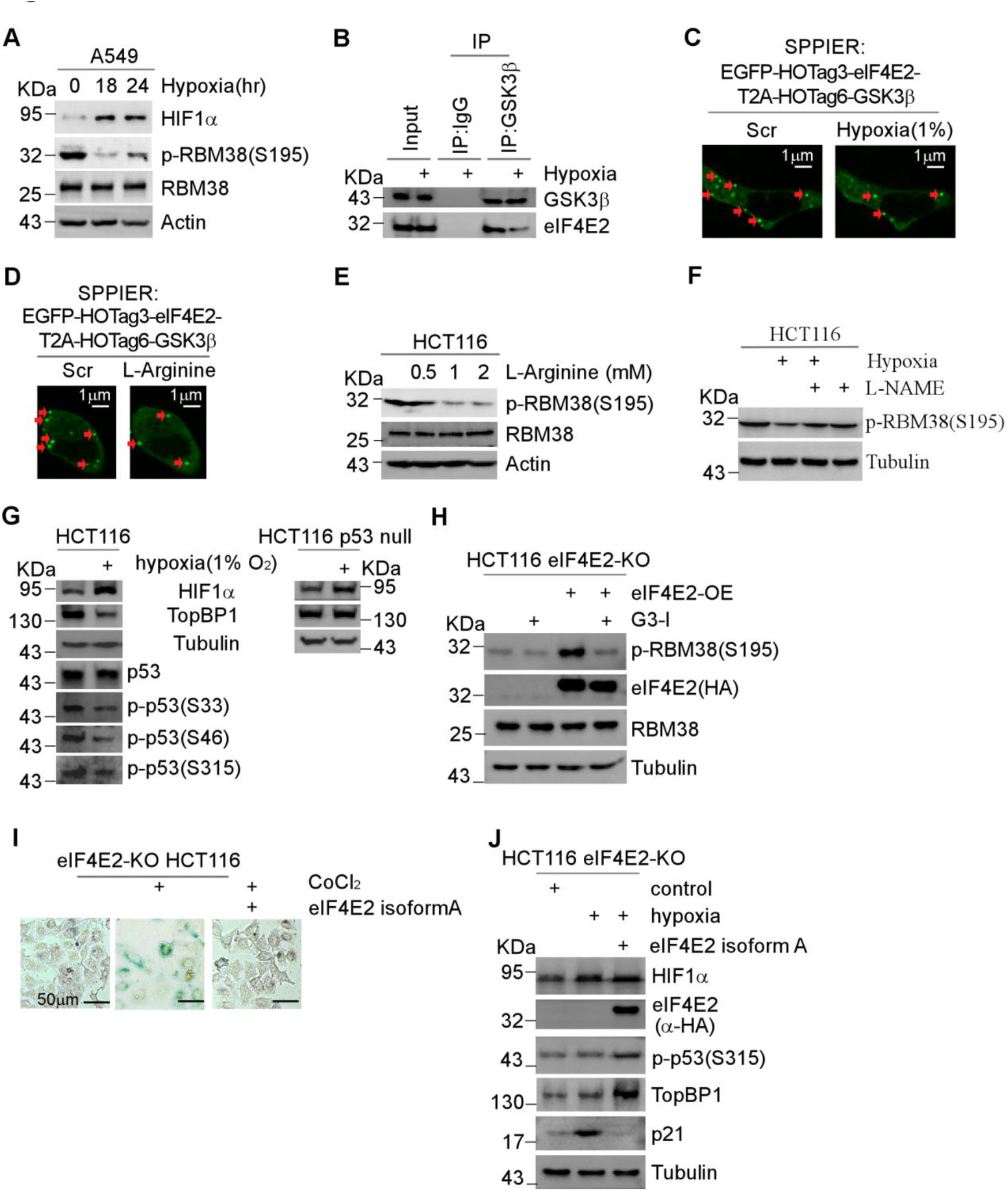
Hypoxia inhibits eIF4E2-GSK3β activity by S-Nitrosylation. **A** Hypoxia inhibits the phosphorylation of RBM38 S195. A549 cells were exposure to hypoxia (1% O_2_) for 18, 24 hours, or normoxia for 24 hour, followed by WB. **B** Hypoxia inhibits eIF4E2-GSK3β binding. Cells were exposure to normal or hypoxia (1% O_2_) condition for 24 hours. Then, cell extracts were subjected to Co-IP with anti-GSK3β antibody or IgG, followed by WB. **C** SPPIER assay showed hypoxia (1% O_**2**_) inhibits eIF4E2-GSK3β interaction. Cells transiently expressed EGFP-eIF4E2-HOTag3-T2A-GSK3β-HOTag6, and then exposure to hypoxia (1% O_2_) for 2 hours. **D** SPPIER assay showed L-Arginine inhibits eIF4E2-GSK3β interaction. The experiment was done as (C) except that 2mM L-Arginine was added to the cells for 1 hour. **E** L-Arginine inhibits the phosphorylation of RBM38 S195. Cells were mock-treated or treated with different concentrations of L-Arginine (0.5, 1, 2mM) for 24 hours and subjected to WB. **F** L-NAME partially restores RBM38 S195 phosphorylation under hypoxia. Cells were mock-treated or treated with 100μM L-NAME under normoxia or hypoxia (1% O_2_) for 24 hours and subjected to WB. **G** Hypoxia inhibits S33, S46, S315 phosphorylation of p53 and the expression of TOPBP1. HCT116 (left) or p53 null HCT116 (right) cells were exposure to normoxia or hypoxia (1% O_2_) condition for 18 hours, followed by WB with indicated antibodies. **H** eIF4E2 isoform A expression activates Rbm38 S195 phosphorylation in eIF4E2-KO HCT116 cells. Vectors expressing eIF4E2 isofom A with HA tag were mock-transfected or transfected into eIF4E2-KO HCT116 cells for 48 hours respectively, along with treatment with 5μM G3-I or scramble peptide as indicated for 24 hours, followed by WB. **I** CoCl_2_ induces senescence in eIF4E2-KO HCT116 cells bypassed by expression of eIF4E2 isoform A. SA-β-Gal staining was performed in eIF4E2-KO HCT116 cells with or without expression of eIF4E2 isoform A, treated with 100 μM CoCl_2_ for 96 hours as indicated. **J** eIF4E2 isoform A expression sustains expression of TOPBP1 or p53 S315 phosphorylation. Vectors expressing eIF4E2 isofom A with HA tag were mock-transfected or transfected into eIF4E2-KO HCT116 cells for 48 hours, followed by WB.

S-Nitrosylation at Cys76, Cys199, and Cys317 inhibits GSK3β proline-directed kinase activity (Wang et al, 2018). Interestingly, Cys317 located in eIF4E2-binding motif of GSK3β. L-Arginine is used as a substrate for the production of nitric oxide to induce S-Nitrosylation (Abat et al, 2008). L-Arginine inhibited eIF4E2-GSK3β interaction demonstrated by the diminished EGFP droplets in SSPIER analysis (Fig 5D). Consequently, L-Arginine inhibited RBM38 Ser195 phosphorylation (Fig 5E). *N*^G^-nitro-L-Arginine methyl ester (L-NAME), a NOS inhibitor, blocks NO generation and inhibits S-Nitrosylation (Wang et al, 1995). Interestingly, L-NAME treatment partially restored Ser195 phosphorylation, which was suppressed by hypoxia (Fig 5F). These results suggested hypoxia inhibits eIF4E2-GSK3β via S-Nitrosylation. Accordingly, hypoxia inhibited S/T-P phosphorylation of p53 including Ser33, Ser46, or Ser315, and inhibited TOPBP1 expression depending on p53 (Fig 5G).

There are 7 isoforms of eIF4E2 in human with different termini, and among them, eIF4E2 isoform A, E, or G have the GSK3β binding motif (Fig EV5B). Knockout cells (eIF4E2-KO HCT116) were generated by CRISPR/Cas9 technology specially targeting isoforms with GSK3β binding motif (Fig EV5C). As expected, reexpression of eIF4E2 isoform A activated RBM38 S195 phosphorylation in eIF4E2-KO cells, which can be blocked by G3-I (Fig 5H). Functionally, we used CoCl_2_ (a mimic of hypoxia) and hypoixa (1%) to induce senescence. A long period of time is needed for inducing senescence. In that period, hypoxia inhibited the growth of eIF4E2-KO HCT116, making senescence difficult to detect. But we observed that eIF4E2 isoform A expression preserved the normal growth of eIF4E2-KO HCT116 under hypoxia (Fig EV5D and E). CoCl_2_, used at low concentration of 100 μM, is also toxic to eIF4E2-KO HCT116 cells. But, there was still a fraction of cells remaining in the alive and adherent state showed increased SA-β-Gal staining (Fig 5I). And expression of eIF4E2 isoform A prevented CoCl_2_-induced senescence (Fig 5I), which increased TOPBP1 expression and p53 S315 phosphorylation in eIF4E2-KO HCT116 cells (Fig 5J).

### Mammalian-unique eIF4E2 protects tissues against hypoxia

eIF4E2 is evolutionary conserved in vertebrates. But startlingly, according to NCBI GenBank, GSK3β binding motif of eIF4E2 only appears in mammals (Fig 6A). We have shown that the effect of inhibiting eIF4E2-GSK3β is not significant in wild-type mice. However, it is interesting that under physiological hypoxic (10% O_2_) conditions, intraperitoneal injection of e2-I strikingly reduced mice viability. After intraperitoneal injection of peptide, mice were housed in normoxic or hypoxic (10% oxygen) conditions respectively. As Fig 6B and C showed, ambulation counts of the e2-I injection group halved after exposure to physiological hypoxia condition, compared with around the 18 meters within 5 min of the scrambled peptide injection group. For comparison, the effect of e2-I injection under normoxic conditions is negligible. Consistent with previous reports (Baitharu et al, 2013), hypoxia decrease the rearing counts of mice (Fig EV6A). And the e2-I injection further decreased the rearing counts of mice to half, compared with scrambled peptide injection (Fig EV6A). Although e2-I did not induce liver senescence under normoxia, it induced liver senescence under hypoxia (Fig 6D and E). Under physiological hypoxic conditions, e2-I inhibited p53 S315 phosphorylation or TOPBP1, TRX1 expression (Fig 6F). And immunohistochemistry (IHC) showed that e2-I induced expression of p53 or p21 in liver under physiological hypoxic conditions (Fig EV6B).

**Figure 6.**
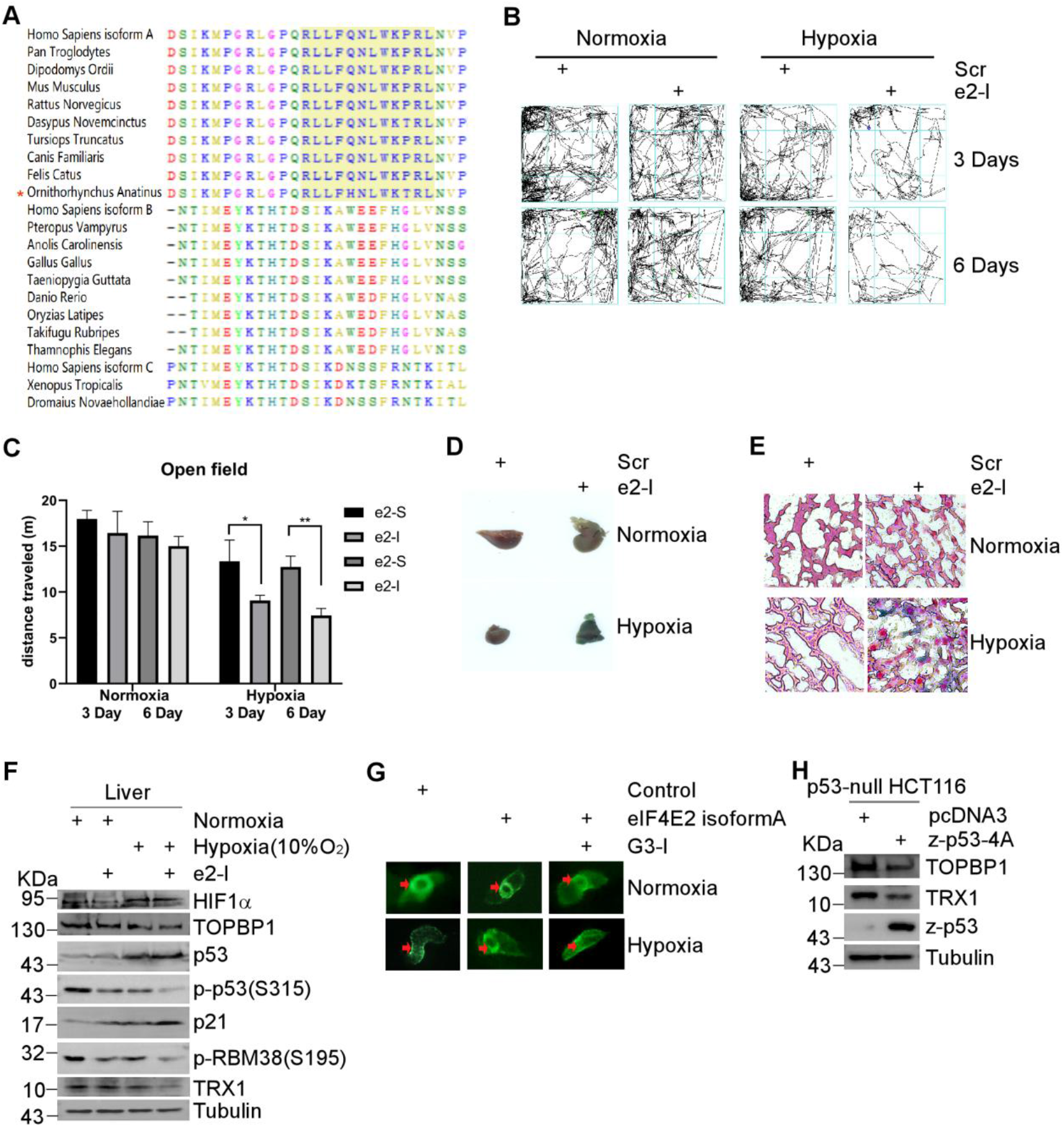
Mammalian-unique eIF4E2 protects tissues against hypoxia. **A** Multiple-sequence alignment of c-termini of eIF4E2 from major species, including mammals, reptiles, amphibians, fish, and birds. **B** Representative tracks of mice in the open field chamber over 5 min. After consecutive intraperitoneal injection of 35mg/kg e2-I or scrambled e2-S every day for 2 days, mice were housed at normaxia or physiological hypoxia (10% O_2_) conditions and tracks of mice were records at 3 or 6 days respectively. **C** Ambulation counts of mice in experiment (B). Each value represents the mean ± SEM of 3 mice. Significance was evaluated with the *t*-test and repeated measures ANOVA. \*\*\**P* <.01. **D** e2-I promotes liver senescence under hypoxia. The experiment was done as (B), and on the 5^th^ day, SA-β-gal staining of liver tissue blocks were performed. **E** The SA-β-gal stained liver tissues shown in (D) were sectioned and then counterstained with nuclear fast red. **F** Equal amounts of liver tissue extracts from experiment (D) were subjected to WB with indicated antibodies. **G** Human eIF4E2 isoform A prevents disorder of heart looping of zebrafish. Representative Tg(cmlc2:eGFP) heart were showed at 48 hours post-fertilization (hpf). Vectors expressing eIF4E2 isoformA were mock-injected or injected into embryos (n = 3) at the 1-cell stage. 24 hours later, embryos were exposure to hypoxic (1 %) or normoxia for 24 hours. **H** Mutant zebrafish-p53-4A suppresses the expression of TOPBP1 or TRX1. Vectors expressing zebrafish-p53-4A, were mock-transfected or transfected into p53-null HCT116 cells for 48 hours before WB.

Genome sequence analysis and cDNA cloning confirmed zebrafish eIF4E2 lacks GSK3β binding motif. Then, human eIF4E2 isoform A was introduced into Tg(cmlc2:eGFP) embryos, which rescued the abnormal heart loop caused by hypoxia stress (Fig 6G). Notably, G3-I hindered this rescue effect (Fig 6G). RT-PCR results showed that eIF4E2 expression increased the expression of TOPBP1 or TRX1 (Fig EV6C). It is hard to prove this rescue depends on p53 at this moment. But interestingly, we observed that consistent to human p53-6A, overexpression of zebrafish p53-4A with S/T-X_0-1_-P mutated to A-X_0-1_-P significantly inhibited expression of TOPBP1 or TRX1 (Fig 6H and Fig EV6D).

## Discussion

Hypoxia modulates senescence in a variety of ways which may be implicated in aging or cancer, but there is still great controversy about the effect of hypoxia on senescence in living organisms (Yeo, 2019). In this study, we revealed a specific eIF4E2-GSK3β pathway in mammals counters senescence and hence preserve tissues against hypoxia by maintaining p53 at multi-S/T-P, which transcriptionally inhibiting TOPBP1 or TRX1.

Surprisingly, eIF4E2, as a translation modulator, interacts and regulates GSK3β proline-directed kinase activity (Fig 1). GSK3β substrates harboring the priming motif (Beurel et al, 2015), but it is neither required nor sufficient to guide GSK3β-dependent phosphorylation (Shinde et al, 2017). GSK3β is also a proline-directed kinase (Hooper et al, 2008), and it is the first time to reveal a specific mechanism targeting this proline-directed kinase activity. Impressively, other proteins (including FRAT, MACF1, and NINEIN) shares this conserved GSK3β-binding motif, which may specifically regulate GSK3β (Fig 1 and Fig EV1) (Freemantle et al, 2002; Hong et al, 2000; Ka et al, 2014). Actually, FRAT proteins is already determined to regulate GSK3β through this conserved motif (Stoothoff et al, 2005; Thomas et al, 1999). More consistently, a study showed that FRAT-GSK3β regulates phosphorylation of Foxk1, and half of four potential phosphorylation sites of Foxk1 regulated by TRAT-GSK3β are proline-directed (T407-P and S428-P) (He et al, 2019; He et al, 2018). Hence, a novel mechanism targeting GSK3β proline-directed kinase activity has been revealed, which may provide clues for the development of more specific GSK3β inhibitors.

Although p53 is a canonical inducer of cellular senescence, it can also suppress senescence (Demidenko et al, 2010). Now we found that basal p53 harboring multi-S/T-P phosphorylation is critical in suppressing senescence and dephosphorylation leads to senescence by inhibiting transcription (Fig 2–4). Normally, phosphorylation positively affects transcriptional activity of p53. Here, dephosphorylation not only curbs p53 transactivation activity, but confers p53 transcriptional repression activity (Fig 3). Indeed, multi-S/T-P dephosphorylation also turns zebrafish p53 into a transrepressor (Fig 6). p53 is structurally and functionally conserved from flies to mammals. For example, p53 of humans and zebrafish are both transrepressor against hypoxia (Feng et al, 2011). Hence, our studies suggest a general mechanism for p53 transcriptional repression activity, which is not rare (Jiang et al, 2015; Li et al, 2019; Wang et al, 2016).

eIF4E2-GSK3β regulates p53 in a dual but opposite way through translation or stability, both of which depend on S/T-P phosphorylation. Impressively, e2-I or G3-I, peptide inhibiting eIF4E2-GSK3β, intensely induced liver senescence in *RBM38^S193A^* mutant mice, where excluding the effect of RBM38 dephosphorylation (Fig 4). The coordinated effects of multiple phosphorylation is robust and will no longer be compromised in vivo, compared to single phosphorylation of p53 (Kruse & Gu, 2009; Lee et al, 2011). Meanwhile, peptide hardly induced liver senescence in wild-type, indicating that of RBM38 S-P dephosphorylation opposes the effect on senescence of p53 S/T-P dephosphorylation. In short, the physiological p53 activity seems to be strongly regulated by the eIF4E2-GSK3β pathway, but it is also very subtle, which is quite necessary if considering pivotal roles of p53 in various physiological contexts (Kastenhuber & Lowe, 2017). Furthermore, the disclosure of the dual effects of eIF4E2-GSK3β leads to effective cancer suppression strategy by combining peptide e2-I with Pep8. e2-I induced-senescence inhibited NSCLC A549 xenograft tumor growth in p53-dependent manners, while peptide Pep8 greatly boosted this effect by negating the negative effect of RBM38 dephosphorylation on p53 (Lucchesi et al, 2019). Especially, the low concentration of Pep8 is impotent, but when combined with e2-I, it is very effective. Pep8 may act as senolytic peptide to eliminate e2-I induced senescent cells by inducing cytoplasm p53 (Baar et al, 2017). Of note, eIF4E2 activation is potential diagnostic marker and therapeutic target for non-small-cell lung cancer (NSCLC) (Uniacke et al, 2014). Further exploring eIF4E2-GSK3β for prevention of NSCLC should be guaranteed.

A recent study showed that S-Nitrosylation at cysteine (Cys) inhibits GSK3β proline-directed kinase activity (Wang et al, 2018). Echoing to them, Cys317 is coincidentally located in the eIF4E2 binding-motif of GSK3β, and S-Nitrosylation mediates hypoxia to inhibit eIF4E2-GSK3β pathway by disrupts their binding (Fig 5). Several lines of evidence have suggested that modification of eIF4E2 plays a role in stress response (Okumura et al, 2007; von Stechow et al, 2015). Considering eIF4E2 is a hypoxia-responsible protein, its modification may mediate hypoxia to regulate GSK3β. It is already known that hypoxia induces p53 protein accumulation in nuclear, allowing it to function as a transrepressor (Feng et al, 2011; Koumenis et al, 2001), and hypoxia inhibits TOPBP1 (Pires et al, 2010). Our studies suggested S/T-P dephosphorylation, due to inhibition of eIF4E2-GSK3β, mediates the effect of hypoxia on p53, which subsequently transcriptionally inhibits TOPBP1.

Notably, this eIF4E2 that interacts with GSK3β only exists in mammals. More important, we found that eIF4E2-GSK3β prevents liver senescence of mice against hypoxia (Fig 6). Hence, intraperitoneal injection of e2-I promotes liver senescence, and severely impact mice viability under physiological hypoxia condition. And interestingly, exogenous mammals eIF4E2-GSK3β protected zebrafish against hypoxia, which is reflected in its ability to preserve heart looping. It is recently reported that eIF4E2 is highly correlated with hypoxia adaption of mammals to Tibetan Plateau (Ouzhuluobu et al, 2020; Zhang et al, 2019a). Specially, a 63bp-insertion, located between exons responsible for the different c-termini, occurs in eIF4E2 of Tibetan (Ouzhuluobu et al, 2020). Inspired by these observations, our studies suggest that eIF4E2-GSK3β pathway is essential for hypoxia adaptation and should be closely related to the resistance of the hypoxia effects on tissue homeostasis.

## Materials and methods

### Reagents and plasmids

The used antibodies, other supplies, the cloning strategy and primers used were listed in supplemental experimental procedures.

### Cell culture, Transfection and RNA Interference

HCT116, p53-null HCT116, eIF4E2-KO HCT116, H1299, MCF7 were cultured in DMEM (Gibco), A549 cells were cultured in RPMI medium 1640 (Hyclone). All media was supplemented with 10% fetal bovine serum (Hyclone), 100 units/ml penicillin, 100 μg/ml streptomycin. All cell lines were cultivated at 37°C in 5% CO_2_ humidity. MEF cells isolation was performed as previously described (Xu, 2005). Briefly, On the 13.5th day of pregnancy, the pregnant female mouse was euthanized via cervical dislocation, and Separate each embryo. Then each embryo was separated from the placenta, membranes, and umbilical cord. The bulk of the CNS tissue was removed by severing the head above the level of the oral cavity and saved for genotyping. Forceps were used to remove the dark red tissue such that the majority of remaining cells were fibroblasts. In a new petri dish, the embryo body was minced in the presence of trypsin. The minced tissue was transferred to a 15 ml conical tube with 5 ml culture media and centrifuged. The supernatant was aspirated and the tissue washed with 1× PBS. The pellet was re-suspended into 2 ml MEFs culture medium (DMEM with 10% FBS, 1% P/S). The embryonic cell and media mixture were then transferred to tissue culture dishes with fresh culture media and placed in 37°C incubator to grow. Plasmid or small interfering RNAs (siRNA) were transfected into cells according to Thermo Fisher protocol (L3000015, Lipofectamine 3000 Reagent). The Small interfering RNAs (siRNA) targeting eIF4E2#1 (5’ CUC ACA CCG ACA GCA UCA A dTdT 3’), siRNA targeting eIF4E2#2 (5’ CAC AGA GCU AUG AAC AGA AUA dTdT 3’) were used. siRNA (5’ UGC GUG UGG AGU AUU UGG AUG dTdT 3’) targeting p53 were used. Scrambled siRNA (5’ UUC UCC GAA CGU GUC ACG U dTdT 3’) were used as control.

### RNA-Seq

HCT116 and p53-null HCT116 cells were treated with 5μM e2-I or e2-S for 24 hours. Each sample group has three biological replicates. A small aliquot of each sample was subjected to western blotting to confirm the inhibitory effect of e2-I on the phosphorylation of RBM38 S195. Stranded paired-end RNA sequencing (RNA-seq) with 100 bp read lengths was performed using a HiSeq2500 (Illumina) according to the manufacturer’s protocol. The acquired sequence reads were aligned to the human genome sequence (hg38) by HISAT2 (version 2.1.0) (Kim et al, 2015). The number of reads per gene was determined using featureCounts v1.6.2 (Scr vs e2-I samples) (Liao et al, 2014). Differentially expressed genes were determined using DEseq (Love et al, 2014). GSEA was performed using the pre-ranked module within the GSEA 4.03 software (Broad Institute) (Subramanian et al, 2005). Genes were ranked by differential expression between scramble peptide and e2-I treated samples and run against C2 KEGG gene sets.

### Phosphoproteomic analysis using iTRAQ LC-MS/MS

HCT116 were treated with 5μM e2-I or the scrambled peptide for 24 hours and whole cell protein lysates were collected. The cell lysates were reduced with 10 mM DTT at 56 °C for 30 min, alkylated with 50 mM iodoacetamide (IAM) at room temperature for 30 min in the dark. The proteolytic solution were subjected to phosphopeptides enrichment using an Immobilized Metal Affinity Chromatography method (Zhou et al, 2013). Then, the peptides were labeled with iTRAQ Reagent-8 plex Multiplex Kit (AB Sciex U.K. Limited) according to the manufacturer’s protocol. Samples were iTRAQ labelled as following: ctrl_1,113; e2I_1, 114; ctrl_2, 115; e2I_2, 116. The enriched phospho-peptides were then dissolved in 2% acetonitrile/0.1% formic acid and analyzed using TripleTOF 5600+ mass spectrometer coupled with the Eksigent nanoLC System (AB SCIEX, USA). The original MS/MS file data were submitted to ProteinPilot Software v4.5 for data analysis. For protein identification, the Paragon algorithm which was integrated into ProteinPilot was employed against uniprot/swissprot-Homo database for database searching (Shilov et al, 2007). And proteins with a fold change larger than 1.5 and p-value less than 0.05 were considered to be significantly differentially expressed. The Motif-x algorithm (http://motifx.med.harvard.edu) was used to extract motifs, and the significance threshold was set to P <1e-6.

### AHA Labeling of Nascent Proteins

AHA Labeling assay was performed as previously described (Zhang et al, 2019b). AHA Labeling of Nascent Proteins Click-iT Metabolic Labeling kits (C10102, C10276, Thermo Fisher) were used for AHA labeling according to manufacturer’s protocol. Briefly, cells were pre-cultured in methionine-free DMEM for 1 hour, then 50 μM AHA was added for 1 hour. Cells were lysed in lysis buffer (50 mM Tris-HCl, pH 8.0, 1% SDS with protease/phosphatase inhibitors). The cleared cell lysate (200 mg protein/sample) was used to perform Click reaction for biotin labeling. Biotinylated proteins were pulled down by streptavidin-agarose beads (50 μL beads/sample, S1638, Sigma) overnight to purify nascent proteins for western blotting analysis.

### The C57BL/6 mouse strain

The RBM38 S193A knockin mice were generated on the C57BL/6 background from Cyagen Biosciences (Guangzhou, China). Animal Experimental Ethics Committee of Huazhong Agricultural University has specifically approved the entire study, the approval # is HZAUMO-2018-014.All the mice were housed in the animal facility at Huazhong Agricultural University. Briefly, C57BL/6 female mice were used as embryo donors and foster mothers, respectively. Superovulated 8-week-old female C57BL/6 mice were mated to C57BL/6 J males, and the fertilized eggs from oviducts were collected. Cas9 mRNA (40 ng/μl), RBM38 gRNA (17.5 ng/μl), and donor oligos (60 ng/μl) were microinjected into both the nuclei and cytoplasm of zygotes for KI mouse production.

### CRISPR/Cas9 Gene Knockout of eIF4E2 in HCT116 Cells

sgRNAs targeting eIF4E2 gene were cloned into pX330-U6-Chimeric_BB-CBh-hSpCas9 (Addgene, #42230) as previously described (Ran et al, 2013). sgRNA sequences are as follows: CAG CCT GCC TGG CAT TCT AGtgg. HCT116 cells were cotransfected with plasmid pX330 and cas9 Nuclease (GenScript, Z03386-50) containing sgRNAs targeting exon 7 of eIF4E2-201. Single-cell clones were selected with puromycin and tested for positive by using PCR, using primers P1: GAG GTC TGC CTC TGG GAC TT /P2: CGA TGA CGC CAG CTT CAA T and three individual clones were used for further experiments.

### Production of adeno-associated viruses

Adeno-associated viruses were generated using packaging plasmids AAV-helper and AAV-8 (Cell Biolabs, Inc) together with pAAV-TOPBP1. Briefy, pAAV-TOPBP1 were co-transfected with AAV-helper and AAV-8 into 293 cells, respectively, by a calcium phosphate-based protocol. Viral particles were harvested following 48h of transfection. After purification by CsCl gradient centrifugation, the titers of viral particles were determined by quantitative real-time PCR and these vectors were applied in the following animal experiments. AAV-TOPBP1(1×10^11^viral genomes in 100μl saline) were intravenous injected into RBM38S193A mutant mice (n=3). 2 weeks later, mice were treated intraperitoneally with 35mg/kg peptide e2-I or scrambled e2-S every two days.

### Open field test

After consecutive intraperitoneal injection of 35mg/kg e2-I or scrambled e2-S every day for 2 days, the mice were housed at normoxia or physiological hypoxia (10% O_2_) conditions. on the 3^th^ or 6^th^ day, the mice were subjected to an open field test. The activity was measured as the total distance traveled (meters) and rearing times in 5 min in the open field chamber (40 cm long × 40 cm wide × 25 cm high). The center square of the open field, comprising 25% of the total area, was defined as the “central area” of the open field. Mice used for the test were 8 weeks old.

### Senescence-associated β-galactosidase (SA-β-Gal) assay, Xenograft

and other methods were described in supplemental experimental procedures.

## DATA AVAILABILITY

The RNA-Seq data: Sequence Read Archive (SRA) PRJNA605921 (https://dataview.ncbi.nlm.nih.gov/object/PRJNA605921)

The iTRAQ LC-MS/MS proteomics data: iProX IPX0002016000 (https://www.iprox.org//page/project.html?id=IPX0002016000).

